# A machine learning model that emulates experts’ decision making in vancomycin initial dose planning

**DOI:** 10.1101/2021.09.16.460731

**Authors:** Tetsuo Matsuzaki, Yoshiaki Kato, Hiroyuki Mizoguchi, Kiyofumi Yamada

**Author notes:** Corresponding author: Kiyofumi Yamada, Hospital Pharmacy, Nagoya University Graduate School of Medicine, Nagoya, Aichi, 466-8560, Japan.

## Abstract

Vancomycin is a glycopeptide antibiotic that has been used primarily in the treatment of methicillin-resistant *Staphylococcus aureus* infections. To enhance its clinical effectiveness and prevent nephrotoxicity, therapeutic drug monitoring (TDM) of trough concentrations is recommended.

Initial vancomycin dosing regimens are determined based on patient characteristics such as age, body weight, and renal function, and dosing strategies to achieve therapeutic concentration windows at initial TDM have been extensively studied. Although numerous dosing nomograms for specific populations have been developed, no comprehensive strategy exists for individually tailoring initial dosing regimens; therefore, decision making regarding initial dosing largely depends on each clinician’s experience and expertise.

In this study, we applied a machine-learning (ML) approach to integrate clinician knowledge into a predictive model for initial vancomycin dosing. A dataset of vancomycin initial dose plans defined by pharmacists experienced in vancomycin TDM (i.e., experts) was used to build the ML model. The target trough concentration was attained at comparable rates with the model- and expert-recommended dosing regimens, suggesting that the ML model successfully incorporated the experts’ knowledge. The predictive model developed here will contribute to improved decision making for initial vancomycin dosing and early attainment of therapeutic windows.

## Introduction

Vancomycin is a glycopeptide antibiotic that has been in clinical use for more than 50 years, primarily for the treatment of methicillin-resistant *Staphylococcus aureus* (MRSA) infections (1). To maximize its clinical effectiveness and avoid nephrotoxicity, therapeutic drug monitoring (TDM) is recommended. The ratio of the vancomycin area under the concentration-time curve (AUC) to the minimum inhibitory concentration (MIC), AUC/MIC, is a primary predictor of vancomycin effectiveness. However, since it is challenging to determine the AUC due to the need to obtain multiple serum vancomycin concentrations, it is recommended to monitor the trough serum concentration, which is considered to be the most accurate and practical surrogate marker for the AUC (1). The trough concentration of vancomycin should be above 10 mg/L and below 20 mg/L to avoid development of resistance and nephrotoxicity, respectively (1–3). Thus, the therapeutic range for vancomycin trough levels is 10–20 mg/L, while several guidelines have recommended vancomycin trough levels of 15–20 mg/L for serious invasive MRSA infections such as sepsis (4, 5).

Initial vancomycin dosing consists of a single loading dose followed by a series of maintenance doses; the usual loading dose and maintenance dose are 25–30 mg/kg and 15– 20 mg/kg, respectively (4). Loading and maintenance doses are adjusted for patient characteristics such as age, gender, body weight (BW), and renal function. In clinical practice, initial dosing is calculated using population pharmacokinetic parameters, and numerous population pharmacokinetic studies of vancomycin have been conducted to develop nomograms designed to achieve therapeutic trough concentrations at initial TDM (6–8). However, because these nomograms were constructed and validated within specific populations, their robustness to population change is limited. To date, there is no comprehensive strategy for individually optimized initial dosing, and vancomycin dosing decisions are often made based on clinicians’ experience and expertise (hereafter, prior knowledge) (7, 9, 10). In a recent survey in the United States and Canada, many healthcare institutions indicated that institutional-level credentialing or training was important to achieve appropriate vancomycin TDM (11). Indeed, several studies demonstrated that initial dose planning by pharmacists engaged in vancomycin TDM could lead to higher target attainment rates, emphasizing the importance of prior knowledge when conducting vancomycin dose planning (12, 13).

Machine learning (ML) algorithms promote the discovery of new techniques and improve decision making on specific questions involving abundant and multi-dimensional data, and they have emerged as a promising approach for medical research and clinical care. Advances in ML have facilitated the discovery of new biomarkers and mutations related to prognosis, and also the development of automated diagnosis tools (14). Recently, ML approaches have also been used in the field of TDM. Imai and colleagues utilized an ML approach to build a nomogram for initial vancomycin dosing using a dataset of patients treated with vancomycin (6). A study by Huang et al. applied variable engineering and ML methods that enabled integration of high-dimensional data to build a predictive model for maintenance dosing (15).

Given the importance of prior knowledge in vancomycin dose planning, here we sought to integrate such knowledge into an ML model for initial vancomycin dosing. Toward this goal, we used a dataset of vancomycin initial dosing regimens that were defined by pharmacists experienced in TDM (hereafter, TDM experts). This straightforward approach yielded a predictive model that reproduces regimens defined by TDM experts. Notably, a therapeutic window was achieved at a similar rate using ML- and expert-recommended dosing regimens, whereas a previously developed ML model failed to predict dosing regimens that attained a therapeutic window (6).

## Materials and methods

### Study subjects

This was a single-center, retrospective, observational study of hospitalized patients who received intravenous vancomycin from May 2017 to May 2021 at Nagoya University Hospital. TDM experts were defined as pharmacists who were either (i) engaged in vancomycin TDM or (ii) capable of conducting vancomycin initial dose planning with similar success as the pharmacists defined in (i). Patients who commenced vancomycin treatment with TDM expert–recommended dosing during the study period were included. The exclusion criteria were as follows: under 18 years of age; undergoing peritoneal dialysis or hemodialysis (including continuous hemodiafiltration); receiving multiple loading doses; diagnosed with amyotrophic lateral sclerosis (ALS); and missing data on gender, age, BW, body mass index (BMI), or serum creatinine.

Thirteen patients received two courses of vancomycin treatment with pharmacist-designed initial dosing regimens. In this group, each regimen was independently designed based on patient characteristics at start of vancomycin treatment, and therefore we included each of these patients as two cases in the study dataset.

### Building of the ML model

The dataset used in this study included clinical and routine laboratory data, initial dosing regimens, and serum vancomycin concentrations at initial TDM (if measured). Age, gender, BW, BMI, and creatinine clearance (CL_CR_) calculated by the Cockcroft-Gault equation were used as parameters to predict initial dosing regimens (including loading and maintenance doses), because we routinely estimated individualized dosing regimens based on these data, as recommended by previous studies (10, 16). The dataset (n = 106) was divided into training and testing datasets in a ratio of 84:22 (approximately 80:20).

We developed an ML model by applying random forest (RF) classification to the dataset (see the schematic diagram in Fig. 1). An RF is an ensemble of classification (decision) trees that are generated by sampling data and features in the training dataset (17). Prediction is based on a simple majority vote by decision trees. A hyperparameter is a parameter that affects how well a model is trained, and it therefore controls the model performance. In the RF technique, hyperparameters include the number of decision trees (n_estimator) and the maximum depth of the trees (max_depth) (18).

**Fig. 1.**
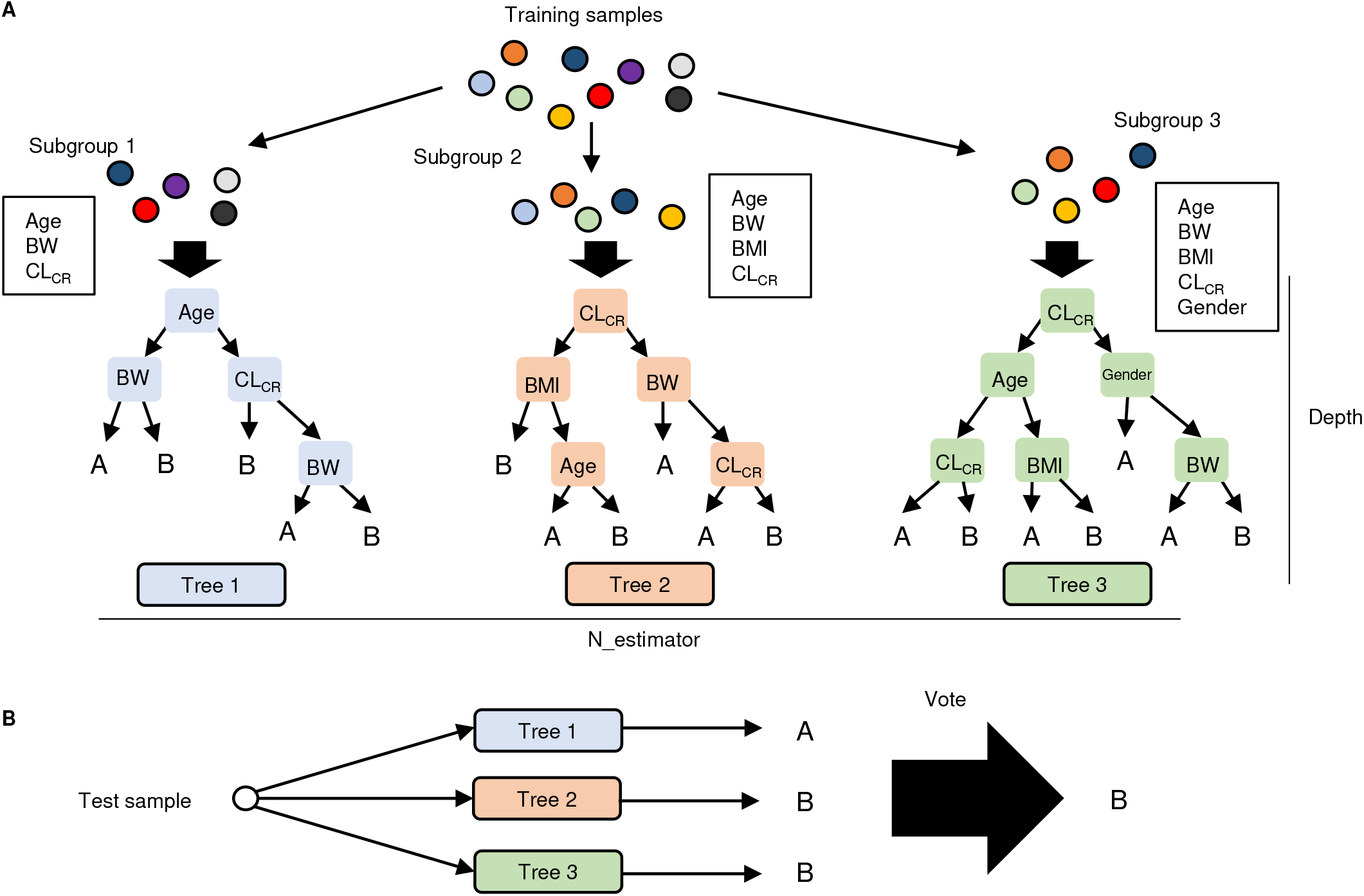
Schematic diagram of RF classification. (A) Training samples and features are randomly sampled; then, classification (decision) trees are constructed with each set of samples and features. The n_estimator is defined as the number of trees. (B) Each decision tree classifies the test sample. The class of the test sample is predicted by majority vote (in this case, the test sample is assigned to class B)

K-fold cross-validation was applied on RF classification to select the most appropriate value of each hyperparameter (n_estimator and max_depth) (19). In this process, the dataset of a training group was further divided into two groups, a training-validation set and a testing-validation set, in the ratio of 67:17 (approximately 80:20) using the five-fold cross-validation method (Fig. 2). The models were then built, and the accuracy scores, which were defined as the ratios of correct predictions to the total number of predictions, were measured in each training-validation/testing-validation set, and the mean accuracy score was recorded. This process was repeated with each pair of hyperparameters (n_estimator: 10, 20, 40, 80, and 160; max_depth: 2, 4, 8, 16, and 32), resulting in optimized hyperparameters that achieved the highest mean accuracy score.

**Fig. 2.**
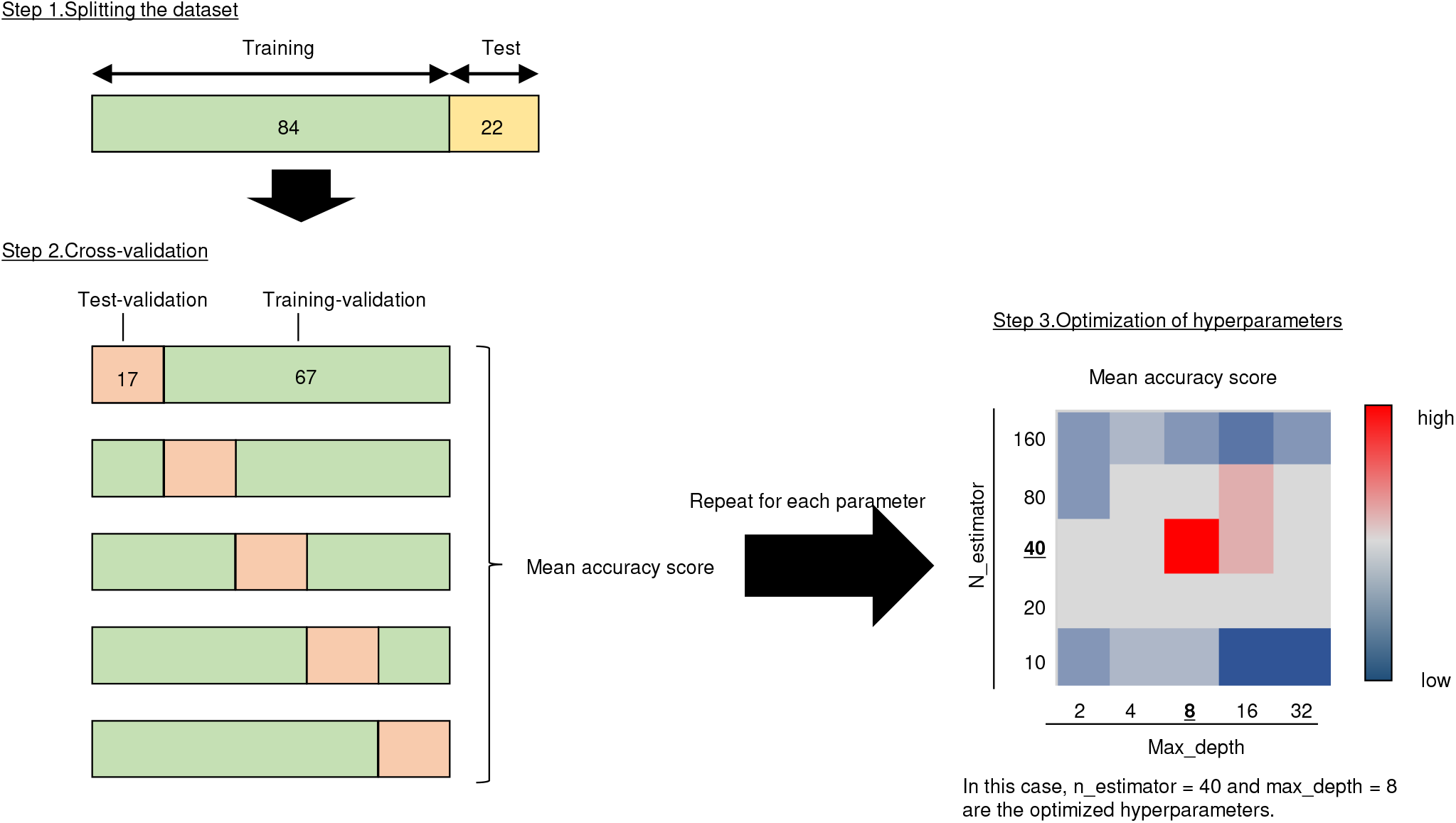
Schematic diagram of steps in the optimization of hyperparameters. Step 1: The dataset is first split into a training set and a testing set in a ratio of 84:22. Step 2: The training set is further split into five partitions, that is, 67:17. Then four partitions and the remaining partition were used as the training and testing sets, respectively, and prediction accuracy was evaluated. The same procedure was repeated for each subset of the dataset, and the mean accuracy score was recorded. Step 3: Step 2 was repeated with different sets of hyperparameters (n_estimators and max_depth), and mean accuracy scores were recorded in each iteration. The set of hyperparameters with the highest mean accuracy scores in Step 2 was obtained as the optimized parameters and used for model construction.

Next, we developed the prediction model with optimized hyperparameters. Feature importance, which represents relative importance in terms of the accuracy of the model, was estimated during training phase (18).

Lastly, we evaluated the performance of the predictive model on the testing data. Accuracy scores for loading doses and maintenance doses were used to evaluate the model.

### Estimation of trough concentrations with the ML model–recommended dosing regimen

Testing data that included serum vancomycin concentrations were used to estimate vancomycin trough concentrations with the ML model–recommended dosing regimen. In this analysis, we excluded cases in which dosing regimens were changed before initial TDM or in which TDM was conducted after just a single loading dose. We also excluded patients who developed vancomycin-associated acute kidney injury (AKI), which was defined as an increase in the serum creatine level of 0.5 mg/dL or a 50% increase from baseline in at least two consecutive measurements, as is consistent with other studies of vancomycin TDM (20, 21). The trough concentration with the ML model–recommended regimen was estimated using the following equation:

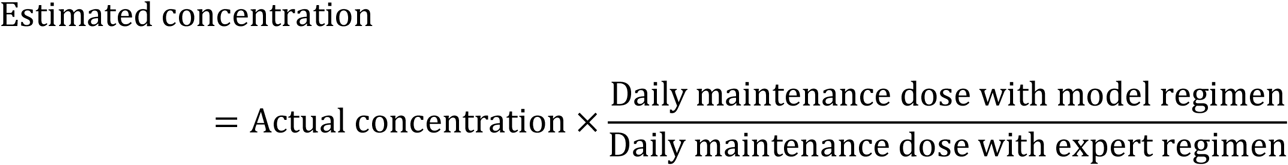

Comparison between the current ML model and the previous ML model reported by Imai and colleagues was performed in patients with a BW ≥ 40 kg and an estimated glomerular filtration rate (eGFR) ≥ 50 mL/min, because the latter model was validated in this population (6).

### Statical analysis

We used the Mann-Whitney test for continuous data, and Fisher’s exact test or the chi-square test for categorical data. All statistical tests were two-tailed, and p values of less than 0.05 were considered statistically significant except in multiple testing (Table 5), where significance was adjusted by Bonferroni correction. Statistical analyses were performed using the Python statistics module.

### Ethics

This study was approved by the ethics committee of Nagoya University Hospital (Approval No. 2021-0189).

## Results

### Patient characteristics

During the study period, there were 140 cases in 127 patients, with cases defined as instances in which initial dosing regimens of vancomycin were determined by TDM experts. Of these, 16 cases were in patients under 18 years of age, 14 were in patients undergoing peritoneal dialysis or hemodialysis, and two were in patients who received multiple loading doses; all of these cases were excluded from this study. One patient whose BMI value was missing and one patient with ALS were also excluded. The remaining 106 cases in 97 patients were included.

The target attainment rate at initial TDM was 71.1% in the total population, which was comparable to the rates reported in another study of expert (pharmacist)-managed vancomycin TDM (64.3%) (12), thus validating our initial dose planning (Table 1). Eligible cases were randomly assigned to a training group and a testing group in the ratio of 84:22 (approximately 80:20). Overall, patient characteristics in the training and testing groups were well balanced (Table 1).

**TABLE 1.** Clinical characteristics of the study subjects

| Characteristics | Total population<br>(n = 106) | Training group<br>(n = 84) | Testing group<br>(n = 22) | P value<br>(training vs<br>testing) |
| --- | --- | --- | --- | --- |
| Age (year) [median (IQR)] | 70.0 (20.3) | 68.5 (21.5) | 73.0 (15.8) | 0.227<sup>a</sup> |
| Gender |  |  |  |  |
| Male [n (%)] | 69 (65.1) | 57 (67.9) | 12 (54.5) | 0.244<sup>b</sup> |
| Female [n (%)] | 37 (34.9) | 27 (32.1) | 10 (45.5) |  |
| BW (kg) [mean ± SD] | 56.3 ± 13.8 | 55.8 ± 13.9 | 58.3 ± 13.5 | 0.391<sup>a</sup> |
| BMI [mean ± SD] | 21.6 ± 5.1 | 21.2 ± 4.7 | 23.1 ± 6.2 | 0.160<sup>a</sup> |
| CL<sub>CR</sub> (mL/min) [mean ± SD] | 85.5 ± 59.2 | 88.5 ± 63.5 | 74.1 ± 37.8 | 0.654<sup>a</sup> |
| Target attainment at initial TDM<br>[achieved/total (%)] | 54/76 (71.1%) | 42/61 (68.9%) | 12/15 (80.0%) | 0.532<sup>c</sup> |

Abbreviations: IQR, interquartile range; BW, body weight; SD, standard deviation; BMI, body mass index; CL<sub>CR</sub>, creatinine clearance; TDM, therapeutic drug monitoring.
<sup>a</sup>Mann-Whitney U test, <sup>b</sup>Chi-square test, <sup>c</sup>Fisher's exact test

### Building an ML model to determine the initial vancomycin dose

Next, we sought to build an ML model for predicting individually tailored vancomycin dosing regimens to achieve the therapeutic window. To this end, we first optimized the hyperparameters for RF classification, then built a predictive model with optimized hyperparameters (Fig. 2). Individual features were ranked based on their relative importance to the accuracy of the model. For loading doses, BW was ranked as the most important feature, followed by CL_CR_, BMI, age, and gender, while for maintenance doses, CL_CR_ was the most important feature, followed by BW, BMI, age, and gender (Fig. 3).

**Fig. 3.**
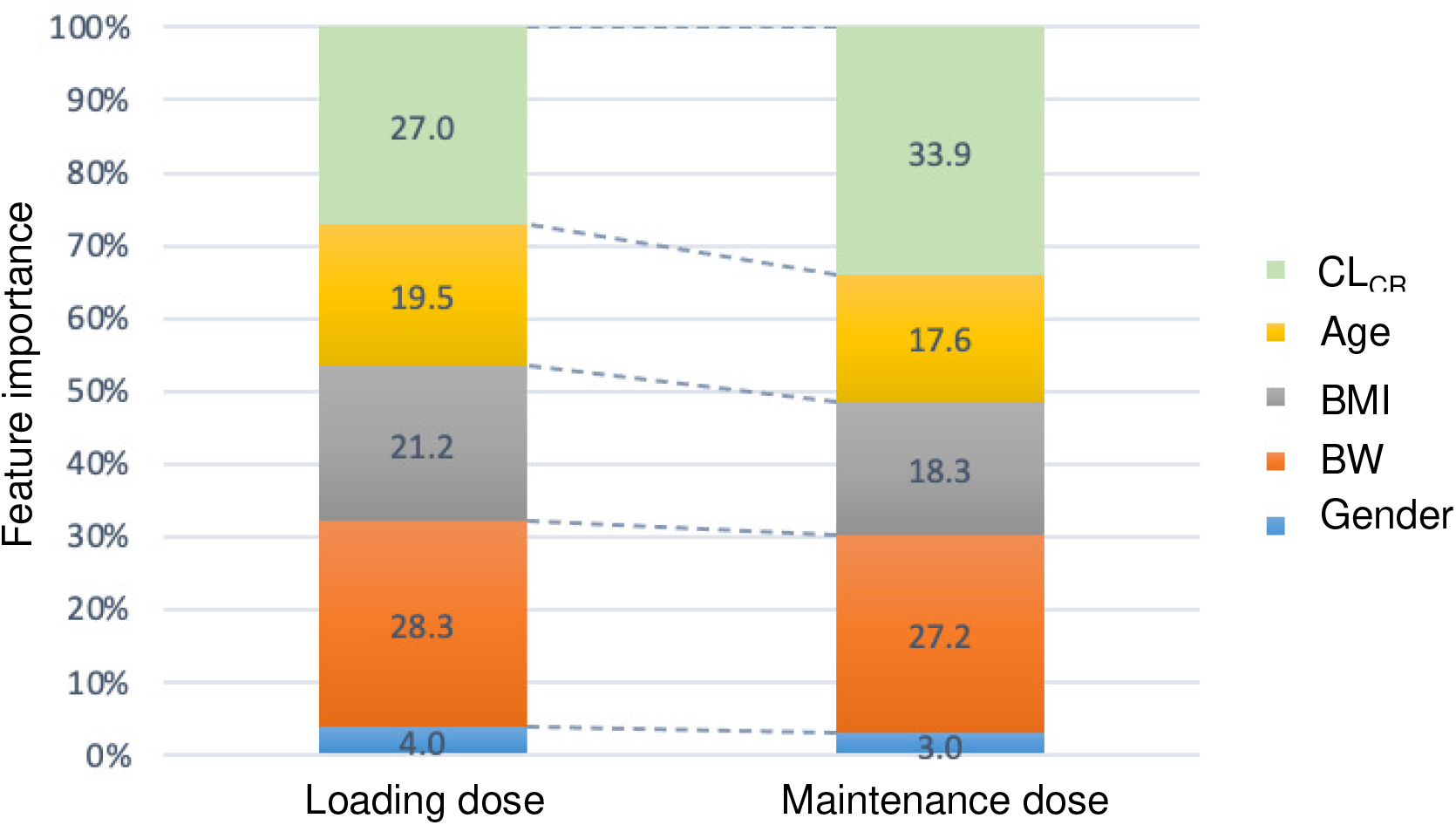
Feature importance in the ML-based prediction model.

### ML model evaluation

Next, we examined whether the ML model generated the same dosing regimen as TDM experts. Table 2 summarizes the performance of the model on the testing dataset. The accuracy scores for loading and maintenance doses were both 63.6% (Table 3). For loading doses, the differences in doses between the expert- and ML model–recommended regimens fell within the range of 50–150%. For maintenance doses, there were two cases in which the differences in doses were larger than two-fold (cases no. 6 and no. 18). The rate of achieving a therapeutic window with ML model–recommended regimens was estimated to be 73.3%, which was comparable to that with expert-recommended regimens (80.0%, p = 1.0; Tables 2 and 4).

**TABLE 2.** Model-recommended dosing regimens and estimated trough concentrations in the testing dataset

| Case no. | Gender | BW<br>(kg) | BMI | Age<br>(year) | CL<sub>CR</sub><br>(mL/min) | eGFR<br>(mL/min) | Loading dose<br>(mg) |  | Maintenance dose |  |  | Trough concentration<br>(mg/L) |  |  | AKI |
| --- | --- | --- | --- | --- | --- | --- | --- | --- | --- | --- | --- | --- | --- | --- | --- |
|  |  |  |  |  |  |  | Expert | Current ML<br>model | Expert<br>(mg/day) | Current ML<br>model<br>(mg/day) | Previous ML model<br>[mg/day (mg/kg/day)]<sup>a</sup> | Expert (actual<br>concentration) | Current ML<br>model<sup>b</sup> | Previous ML<br>model<sup>b,c</sup> |  |
| 1 | Male | 46.0 | 21.3 | 76 | 60.1 | 67.3 | 1000 | 1000 | 1500 | 1500 | 1297 (28.2) | 10.77 | 10.77 | 9.31 | No |
| 2 | Female | 52.6 | 20.5 | 70 | 71.3 | 64.5 | 1000 | 1500 | 2000 | 1500 | 1483 (28.2) | 12.76 | 9.57 | 9.46 | No |
| 3 | Female | 62.0 | 26.0 | 77 | 69.9 | 60.2 | 1500 | 1000 | 2000 | 2000 | 1748 (28.2) | 11.92 | 11.92 | 10.42 | No |
| 4 | Male | 58.3 | 20.0 | 38 | 103.2 | 84.7 | 1250 | 1500 | 2000 | 2000 | ND | NA | ND | ND | No |
| 5 | Female | 49.3 | 24.7 | 88 | 32.9 | 34.2 | 1000 | 1000 | 500 | 500 | ND | NA | ND | ND | No |
| 6 | Male | 54.2 | 20.7 | 82 | 21.4 | 22.7 | 1000 | 1000 | 500 | 1000 | ND | 19.80 | ND | ND | AKI |
| 7 | Male | 58.9 | 22.2 | 80 | 75.4 | 83.2 | 1000 | 1500 | 2000 | 2000 | 1630 (27.7) | 11.70 | 11.70 | 9.54 | No |
| 8 | Male | 47.5 | 16.4 | 78 | 56.0 | 69.7 | 1000 | 1000 | 1000 | 1000 | 1340 (28.2) | 19.00 | 19.00 | 25.45 | No |
| 9 | Male | 58.8 | 22.5 | 82 | 20.3 | 20.3 | 1000 | 1000 | 400 | 500 | ND | NA | ND | ND | No |
| 10 | Female | 48.3 | 19.7 | 79 | 30.5 | 29.8 | 1000 | 1000 | 750 | 750 | ND | 16.91 | 16.91 | ND | No |
| 11 | Male | 65.2 | 21.4 | 65 | 97.0 | 89.5 | 1500 | 1500 | 2000 | 2000 | ND | NA | ND | ND | No |
| 12 | Male | 69.5 | 24.5 | 71 | 96.5 | 88.7 | 1500 | 1500 | 2000 | 2000 | 2537 (36.5) | 7.19 | 7.19 | 9.12 | No |
| 13 | Male | 55.7 | 21.4 | 47 | 71.2 | 58.1 | 1500 | 1500 | 2000 | 1500 | ND | NA | ND | ND | No |
| 14 | Female | 103.0 | 46.4 | 43 | 190.2 | 92.1 | 2000 | 2000 | 3000 | 2000 | 3729 (36.2) | 17.60 | 11.73 | 21.87 | No |
| 15 | Female | 71.0 | 29.2 | 72 | 74.0 | 55.3 | 1000 | 1000 | 2000 | 2000 | 1420 (20.0) | 15.42 | 15.42 | 10.95 | No |
| 16 | Male | 66.0 | 23.7 | 74 | 68.0 | 64.5 | 1500 | 1000 | 2000 | 2000 | 1861 (28.2) | 13.52 | 13.52 | 12.58 | No |
| 17 | Female | 37.1 | 17.2 | 75 | 73.0 | 83.6 | 1000 | 1000 | 1500 | 1500 | ND | 16.37 | 16.37 | ND | No |
| 18 | Female | 45.5 | 17.1 | 63 | 137.9 | 137.7 | 1000 | 1250 | 750 | 1500 | ND | NA | ND | ND | No |
| 19 | Female | 63.5 | 26.1 | 61 | 56.4 | 39.4 | 1500 | 1000 | 1500 | 2000 | ND | 25.27 | 33.69 | ND | No |
| 20 | Male | 48.2 | 18.8 | 68 | 68.9 | 72.9 | 1250 | 750 | 2000 | 1500 | 1759 (36.5) | 13.70 | 10.28 | 12.05 | No |
| 21 | Female | 70.3 | 28.1 | 46 | 90.7 | 56.1 | 1500 | 1500 | 2000 | 2000 | 1863 (26.5) | 33.00 | 33.00 | 30.74 | No |
| 22 | Male | 51.0 | 19.4 | 77 | 65.6 | 75.1 | 1000 | 1000 | 1500 | 1500 | 1413 (27.7) | 10.03 | 10.03 | 9.45 | No |

Abbreviations: BW, body weight; BMI, body mass index; CL<sub>CR</sub>, creatinine clearance; eGFR, estimated glomerular filtration rate; ML, machine learning; ND, data not determined; NA, data not available; AKI, acute kidney injury.
<sup>a</sup>Dose per body weight (mg/kg/day) is also described, because a previous ML model predicted doses in this format.
<sup>b</sup>Concentrations were not determined if patients developed AKI before TDM.
<sup>c</sup>Patients with a BW ≥ 40 kg and an eGFR ≥ 50 mL/min were included.

**TABLE 3.** Relative ML-recommended doses

| Ratio of ML- to expert-recommended dose | Loading dose<br>(n = 22) | Maintenance dose<br>(n = 22) |
| --- | --- | --- |
| > 150% [n (%)] | 0 (0.0) | 2 (9.1) |
| > 125% and ≤ 150% [n (%)] | 2 (9.1) | 1 (4.5) |
| > 100% and ≤ 125% [n (%)] | 2 (9.1) | 1 (4.5) |
| 100% (identical) | 14 (63.6) | 14 (63.6) |
| ≥ 75% and < 100% [n (%)] | 0 (0.0) | 3 (13.6) |
| ≥ 50% and < 75% [n (%)] | 4 (18.2) | 1 (4.5) |
| < 50% [n (%)] | 0 (0.0) | 0 (0.0) |

Abbreviation: ML, machine learning.

**TABLE 4.**
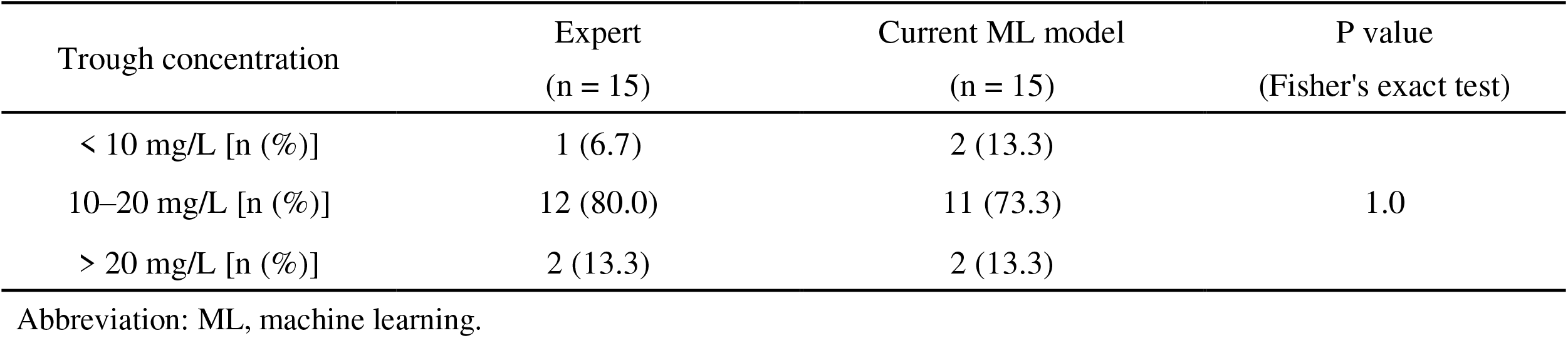
Rates of achieving targeted trough levels with the expert- and current ML model–driven dosing regimens

Lastly, we compared the target attainment rates of our predictive ML model and the ML model developed by Imai and colleagues (6). Because the latter model was validated in patients with a BW ≥ 40 kg and eGFR ≥ 50 mL/min, we compared cases that met these criteria. In this group, the target attainment rates of the expert- and ML model– recommended regimens were 83.3% and 75.0%, respectively, again with no significant difference (p = 1.0; Tables 2 and 5). Importantly, the target attainment rate of regimens recommended by the previous ML model was much lower, at only 33.3%; this indicates the poor ability of that model to predict appropriate dosing regimens, at least using the current testing dataset. However, it should be noted that the differences in target attainment rates compared to our expert- and ML model–recommended regimens did not reach statistical significance (p = 0.0361 and 0.0995, respectively. Note that significance was set at 0.0167 after Bonferroni correction).

**TABLE 5.**
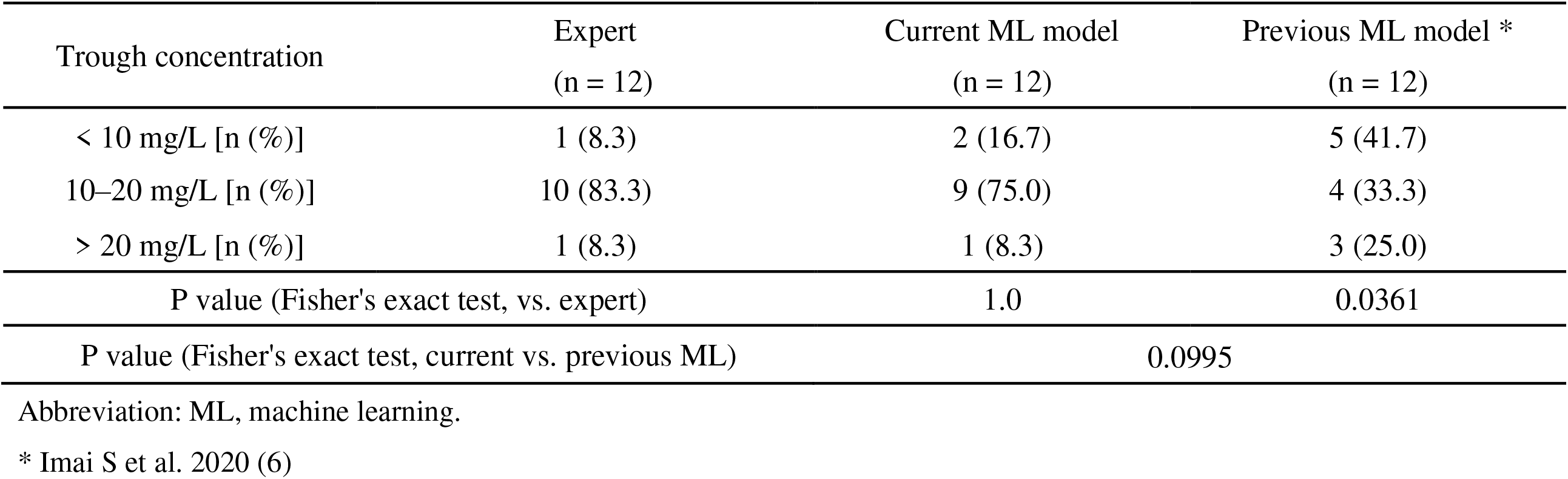
Rates of achieving targeted trough levels with the current and previous ML model–driven dosing regimens

| Trough concentration | Expert<br>(n = 12) | Current ML model<br>(n = 12) | Previous ML model *<br>(n = 12) |
| --- | --- | --- | --- |
| < 10 mg/L [n (%)] | 1 (8.3) | 2 (16.7) | 5 (41.7) |
| 10–20 mg/L [n (%)] | 10 (83.3) | 9 (75.0) | 4 (33.3) |
| > 20 mg/L [n (%)] | 1 (8.3) | 1 (8.3) | 3 (25.0) |
| P value (Fisher's exact test, vs. expert) |  | 1.0 | 0.0361 |
| P value (Fisher's exact test, current vs. previous ML) |  | 0.0995 |  |

Abbreviation: ML, machine learning.
\* Imai S et al. 2020 (6)

## Discussion

Although considerable effort has been made to develop population-specific dosing nomograms for vancomycin, strategies to determine individually optimized initial dosing regimens remain controversial. In practice, decision making about initial dose planning depends on each clinician’s experience regarding the clinical assessment of renal function and the determination of which dosing nomograms to use.

Here, we sought to build an ML model that emulates experts’ decision making concerning initial vancomycin dosing. Toward this end, subjects were limited to those who received an initial dosing regimen defined by TDM experts. In contrast with the previous study of Imai et al. (6), we did not exclude patients with a low BW (< 40 kg) or renal dysfunction (eGFR < 50 mL/min) due to the prevalence of such patients and our desire to assess the robustness of the model to changes in patient characteristics.

Feature importance was largely consistent with predictive covariates conventionally used to determine appropriate dosing regimens: BW and CL_CR_ were the most important features for loading and maintenance doses, respectively (10, 22). Although loading doses are generally determined based on actual BW, our results indicate that CL_CR_ is also a key feature for predicting loading dose (23). This reflects adjustment of loading doses based on renal function, which has been proposed in recent studies (7, 10).

Our ML model scored 63.6% in testing accuracy for both loading and maintenance doses. These modest accuracy scores may have been due to the difficulty of multi-class classification tasks (24); in the current dataset (n = 106), there were six and nine classes (doses) of loading and maintenance doses, respectively (data not shown). In addition, the fact that this study had a relatively small dataset (n = 106), which generally leads to overfitting, may have further contributed to a decrease in predictive accuracy (25).

The target attainment rates were similar for our expert- and ML model–recommended regimens (80.0% and 73.3%, respectively; Table 4). This result indicates that our ML model can competently predict individually tailored vancomycin dosing regimens and thus achieve a therapeutic window. Importantly, the target attainment rate of the ML model developed by Imai et al. was 33.3% (6), which is much lower than that of our model (Table 5). This implies that our model has better predictability, but due to the small sample sizes in this study, the differences in target attainment rates between the two models did not reach statistical significance (p = 0.0995). The discrepant predictive accuracy of the two models is largely attributed to the differences in the training datasets. Differences in ML algorithms also likely played a role; the decision tree algorithm used in Imai et al. is a classification tree (a nomogram), which is easy to comprehend but is more susceptible to overfitting than the RF algorithm used in the present study (6, 17).

The limitations of this study are as follows. First, TDM results were lacking in many study subjects due to the discontinuation of vancomycin treatment before TDM (mainly due to de-escalation). This hindered the evaluation of ML model–recommended dosing regimens regarding their ability to achieve therapeutic levels. Second, the majority of study subjects were older adults, which may have influenced the model’s predictive performance in younger people. Third, our ML model was derived from a single-center, observational study, potentially limiting external generalizability. In addition, it should be noted that the above results are at least partially dependent on the current testing dataset, and thus further evaluation in other populations is required. Lastly and most importantly, the dosing regimens recorded in the dataset were designed to achieve target trough concentrations rather than target AUC/MIC values, raising the concern that our model tends to predict trough-guided dosing (that is, adjusting doses by considering trough concentrations only). This is thought to increase the occurrence of vancomycin-associated AKI compared to AUC-guided dosing, and recently published guidelines do not recommend this approach in patients with serious, invasive MRSA infections (1, 26). A dataset of AUC-guided dosing regimens is needed to build a prediction model for AUC-guided dosing.

Collectively, we developed a novel ML model to predict individually tailored vancomycin initial dosing regimens, and this model had a high rate of achieving therapeutic concentration windows. Our predictive model will aid decision making for initial vancomycin dosing and contribute to early attainment of a therapeutic range, which is crucial for clinical and microbiological success in treating serious infections due to MRSA.

## Acknowledgments

This work was supported by JSPS KAKENHI (Grant Numbers JP 20H03428 and JP19H03532).

